## Supplementary Information for "Open-source personal pipetting robots with live-cell incubation and microscopy compatibility"

### Supplementary Material
#### Supplementary Figures

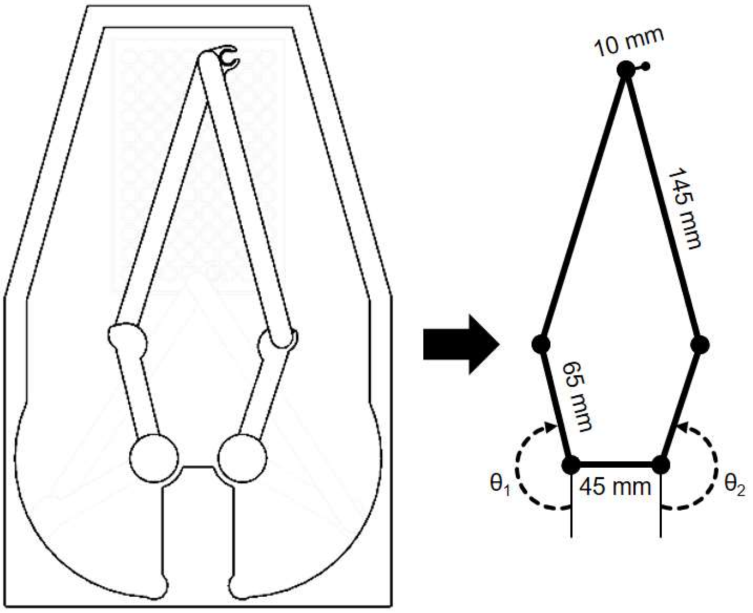


**Supplementary Figure 1: 5-bar planar geometry allows accurate positioning of pipet tips.**
Rotation of 2 stepper motors ($\theta_1$ & $\theta_2$) translates to a specific position in the horizontal plane via 2 freely rotating linkages.

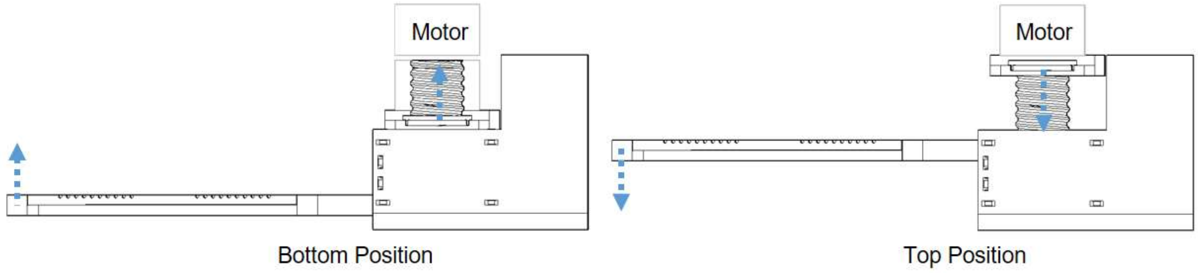


**Supplementary Figure 2: Arm height is controlled by a 3D printed screw.**
Rotation of 2 steppers motors moves the arm platform along the vertical axis.

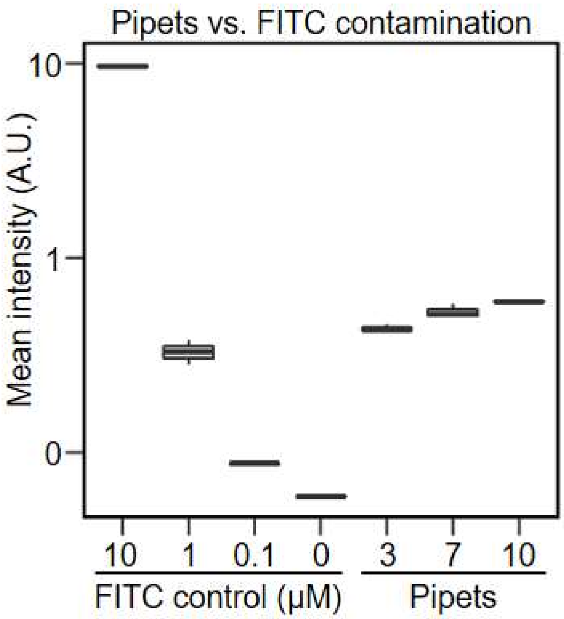


**Supplementary Figure 3: Cross contamination between wells increases with number of pipets used.** To quantify how the number of reusable pipets installed in PHIL affects cross contamination between wells by liquid carry-over, pipet bundles containing 3, 7, and 10 pipets were tested. They were first lowered into 96 well plate wells with 100 µM FITC and then moved into wells with 100 µL PBS. No liquid was pumped, only FITC carry-over by liquid sticking to the pipets' outsides was quantified. Mean fluorescence intensities resulting from FITC carry-over were quantified (right) and compared to a manually pipetted FITC dilution curve (left) in order to estimate FITC carry-over volumes. Carry-over volumes with 3, 7, and 10 pipets in the bundle (right) were approximately 1, 1.5, and 2 µL respectively (n = 3 experiments).

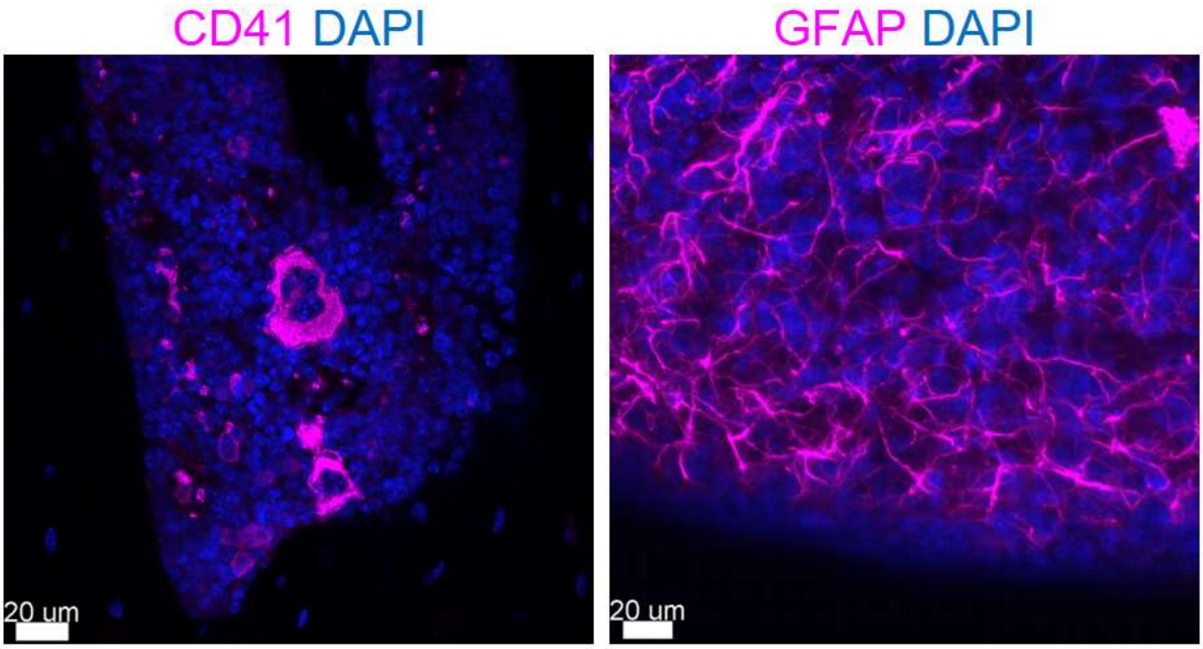


**Supplementary Figure 4: Manually stained bone and cerebrum slices.**
Bone (left) and cerebrum (right) sections with manual immunostaining.

#### Supplementary Movies

**Supplementary Movie 1: PHIL pipetting in a 96 well plate.**

PHIL utilizes 5-bar planar geometry to move pipet tips from well to well in a 96 well plate and dispenses food dye into targeted wells.

**Supplementary Movie 2: PHIL assembly in 2.5 h.**

**Supplementary Movie 3: PHIL pipetting of a 20 s pulse.**

100 µL PBS is replaced with 100 µL FITC before being replaced with 100 µL PBS. Imaged every 0.5 s (mm:ss timestamp).

**Supplementary Movie 4: PHIL pipetting of sequential 15 s pulses to generate oscillations.**

To generate oscillating FITC stimulation patterns, the all 100 µL of a 96 well plate well are sequentially changed from PBS to FITC to PBS every 15 s. Imaged every 0.5 s (mm:ss timestamp).

**Supplementary Movie 5: PHIL pipetting of 2 h oscillations.**

100 µL PBS and FITC oscillations in a 96 well plate are maintained over 48 h. Imaged every 10 min (hh:mm timestamp).

**Supplementary Movie 6: GMPs are unaffected by fluid flow.**

GMPs were automatically washed with 50 µL/s flow. No cell displacement was observed. Imaged every 9 min (hh:mm timestamp).

#### Supplementary Materials and Methods

##### 1. PHIL Assembly Instructions

**!!! The latest version of the assembly instructions can be found on: [https://bsse.ethz.ch/csd/Hardware/PHILrobot.html](https://bsse.ethz.ch/csd/Hardware/PHILrobot.html)**

###### a. Main Body

###### i. Assemble Limit Switches

1. Solder 15cm long red and black wires to 4 limit switches as shown below (See Appendix 1: Basic Soldering).

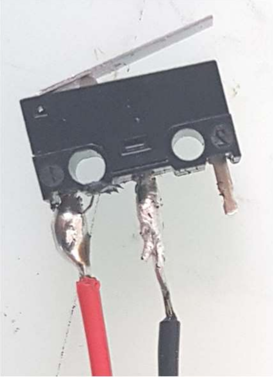

2. Place 1cm long x 2mm diameter heat shrink tubing over soldered joints.
3. Apply air gun at 300°C to all heat shrink segments for 5-6s as shown below.
   a. **Safety Note: The airgun tip is extremely hot. Do not touch it with unprotected skin or place it on combustible objects.**

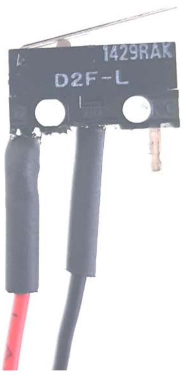


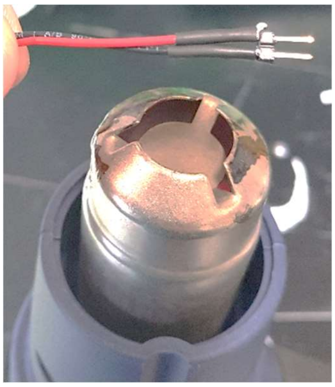

4. Place 1cm long x 2mm diameter heat shrink tubing over each wire.
5. Solder 15cm long red and black wires to 4 male headers as shown below.

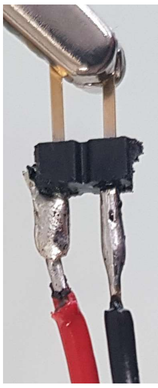

6. Move 1cm long x 2mm diameter heat shrink tubing over each joint and shrink with air gun.

**ii. Assemble Arm Platform**

**a. Supplies:**

**i.** M3 nut x 28
**ii.** 8mm M3 screw x 20
**iii.** 10mm M3 screw x 7
**iv.** 16mm M3 screw x 21
**v.** 8mm bearing x 3
**vi.** 10mm M1.5 screw x 8

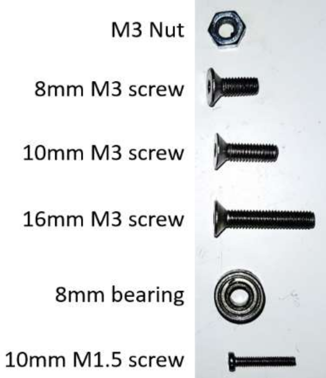


2. Insert 1 M3 nut into the vertical cavity of the Arm columns as shown below.

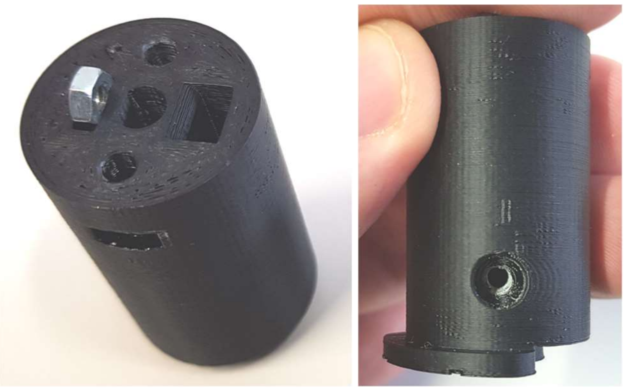

3. Push 1 limit switch connector through each Arm Column as shown below.

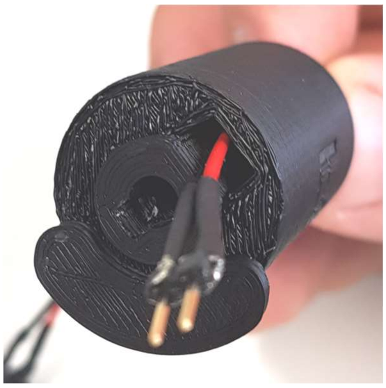

4. Insert 1 stepper motor shaft in each Arm Column and attach securely using 4x 8mm M3 screws as shown below.
   a. Note: Stepper motor shafts have a flat face which aligns with the square cut in the Arm Column sleeve. Some force may be required.

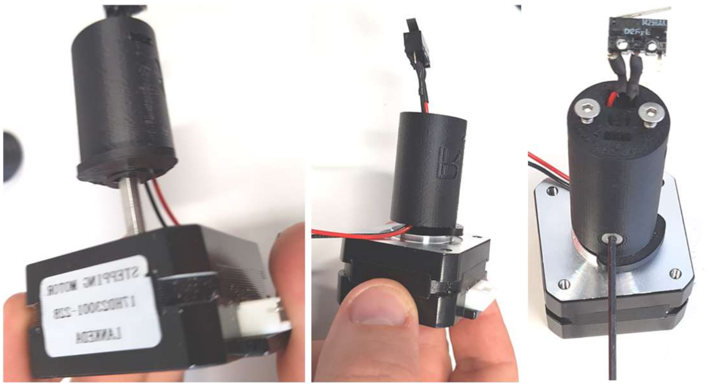

5. Attach each stepper motor to the Arm Platform using 4x 8mm M3 screws as shown below.
   a. Note: Each motor receives its own limit switch.
   b. Note: Align the back of each stepper motor with the back of the Arm Platform as seen in the photo.

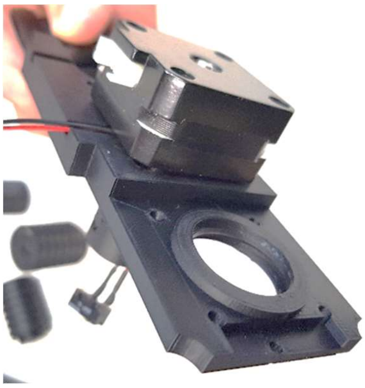

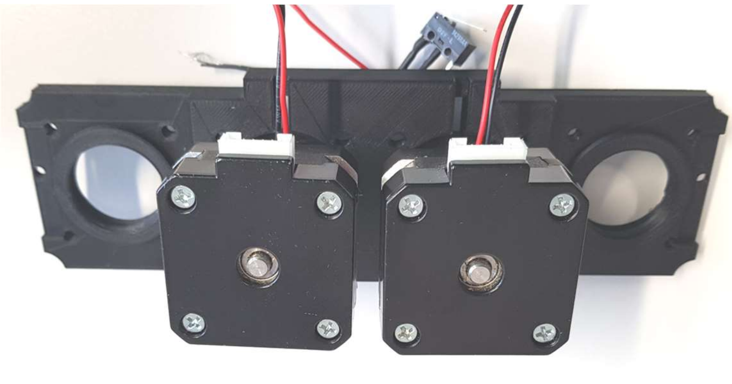

##### iii. Install Z-axis Limit Switches

1. Insert 1 limit switch in each Z-axis slot and fix in place with 2x 10mm M1.5 screws as shown below.
   a. Note: Each motor receives its own limit switch.
   b. Note: Some limit switch movement may be required to align the screws with the mounting holes in the limit switch

bodies. Simply move the limit switches up and down until each screw turns easily.

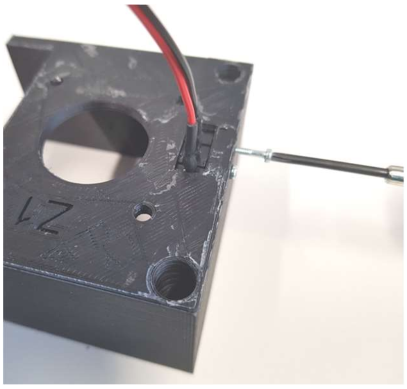



**iv. Install Z-axis Stepper Motors**

1. Insert 1 stepper motor in each side of the Z-axis mounting bracket as shown below.
    a. Note: Align the back of each stepper motor with the back of the Z-axis bracket as seen in the photo.

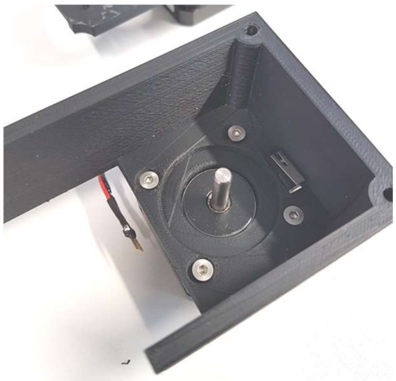

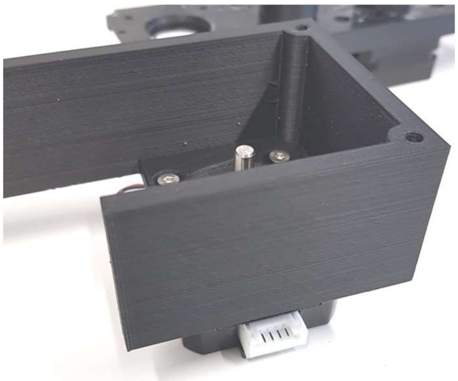

2. Attach each stepper motor to the Z-axis bracket using 4x 8mm M3 screws.
3. Insert 1 M3 nut into the vertical cavity of each Lead Screw as shown below.

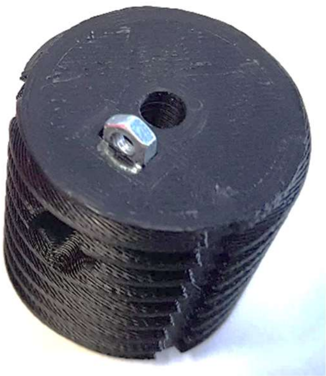

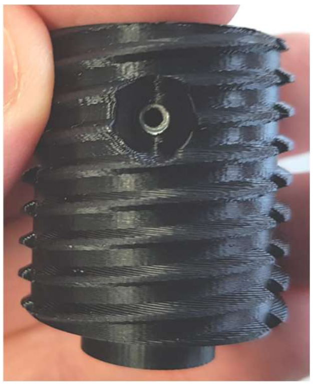

4. Insert 1 stepper motor shaft on each Lead Screw and attach securely using 1x 8mm M3 screws as shown below.

a. Note: Stepper motor shafts have a flat face which aligns with the square cut in the Lead Screw sleeve. Some force may be required.

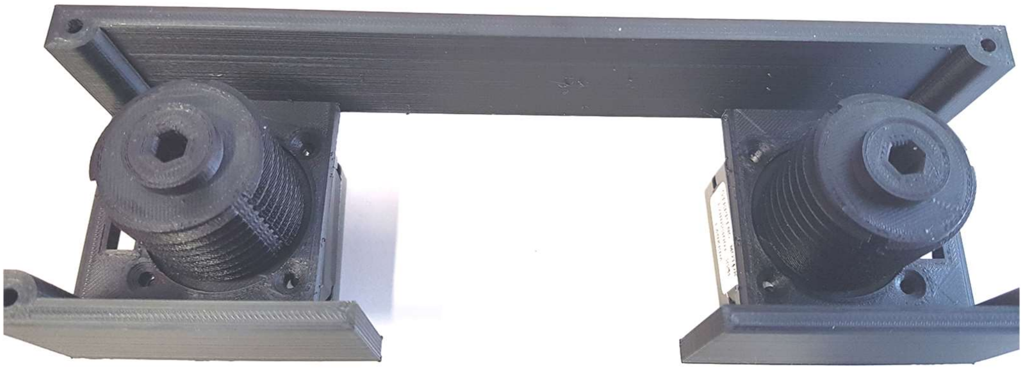

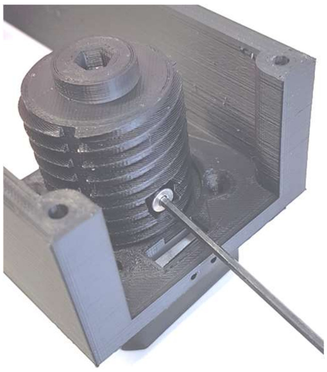

**v. Assemble the main body.**

1. Place the assembled Arm Platform in the Z-axis bracket and simultaneously rotate the Lead Screws until both lead screws protrude ~30m from the Arm Platform.

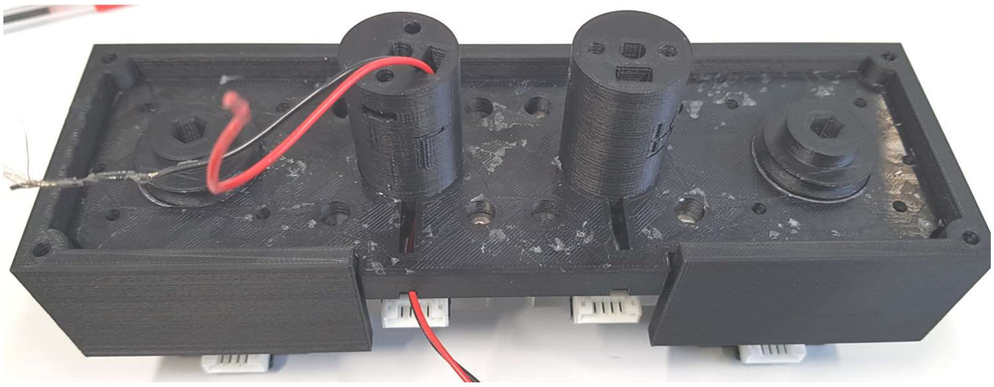

2. Install the assembled Arm Platform and Z-axis bracket on the Robot housing as shown below.

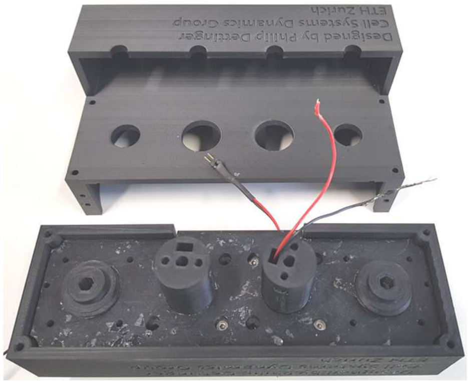

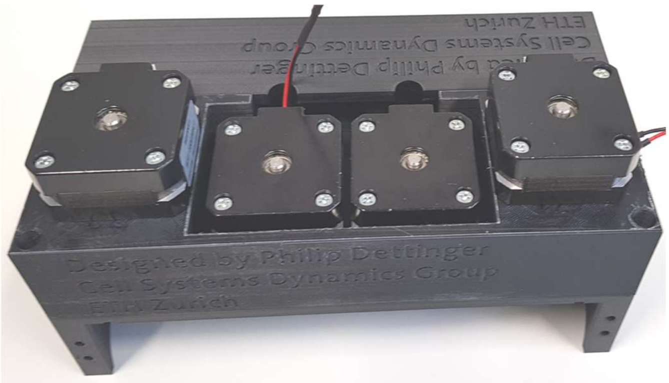

3. Insert 1 M3 nut in each cavity of the main body as shown below.

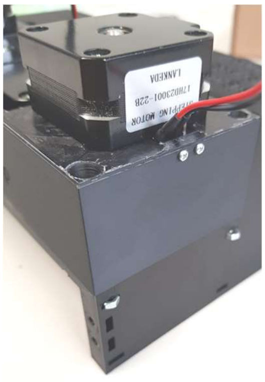

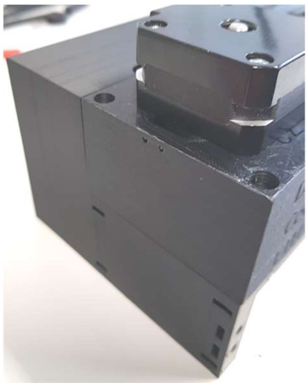

4. Fix the assembled Arm Platform and Z-axis bracket to the main body using 4x 16mm M3 screws as shown below.

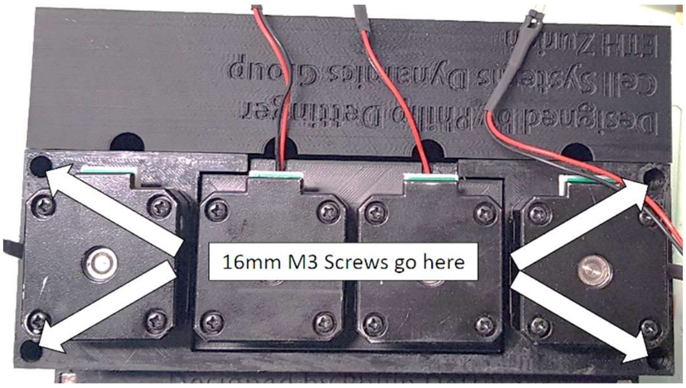


##### vi. Install Left and Right Arms

1. Align the limit switch mounting holes with the corresponding holes in the left and right Arm pivots and fix in place with 2x 10mm M1.5 screws as shown below.
    a. Note: Play close attention to the orientation of the limit switch. The metal lever should point away from the circular base and the slot for the wires depicted in the photo below.

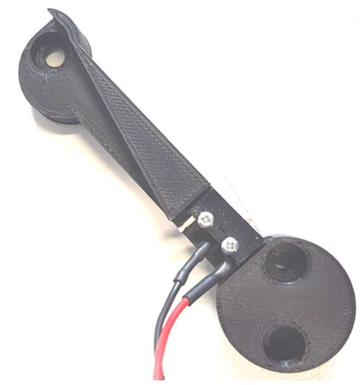

2. Insert 1 M3 nut in each cavity of the Arm columns as shown below.

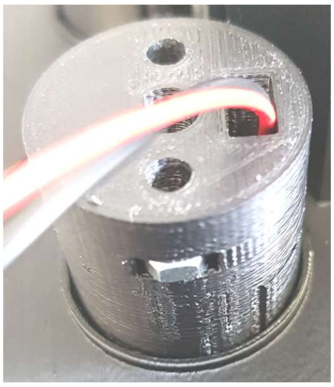

3. Loosely fix the Arm columns to the Arm pivot using 2x 16mm M3 screws as shown below.
    a. Note: Each component is labelled with a letter for its corresponding side. ie L for left and R for right.

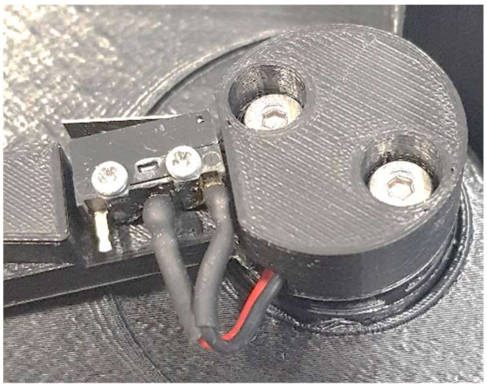

4. Align limit switch wires with the wire channel before fully tightening the Arm pivot screws as shown below.

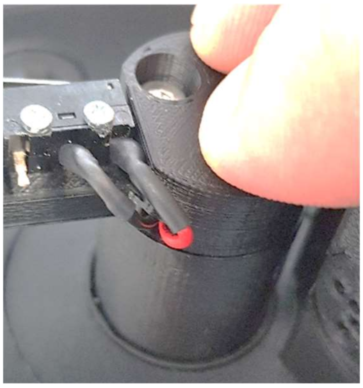

5. Gently pull the limit switch wires through the assembled Arm column and Arm pivot until tight as shown below.

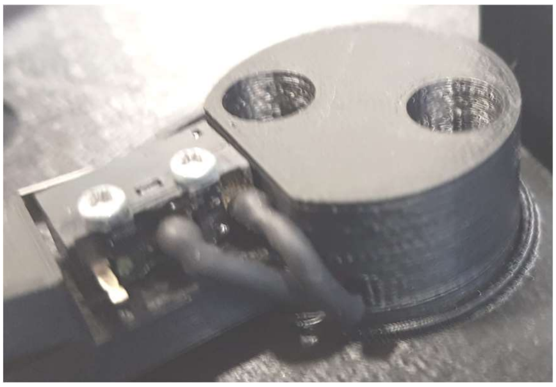

i. **Install Manipulators**

6. Place 1x M3 nut in each hexagonal hole of the left and right Manipulator.
7. Place 1x 8mm bearing in each circular hole of the left and right Arm pivots as well as the right Manipulator as shown below.

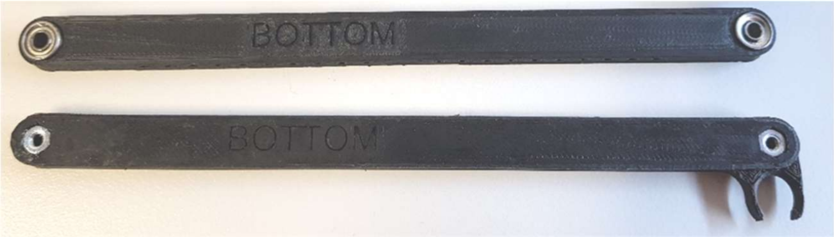


8. Fix the left and right manipulators to the corresponding Arm pivot using 1x 10mm M3 screws as shown below.

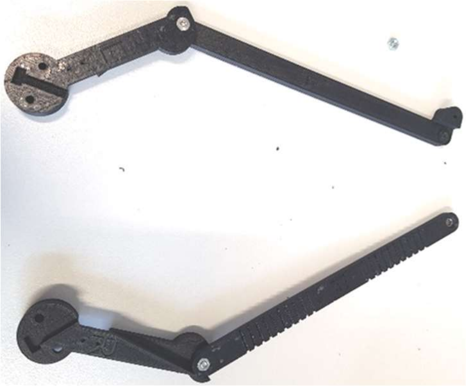

9. Fix the left and right manipulators to each other using 1x 10mm M3 screws as shown below.

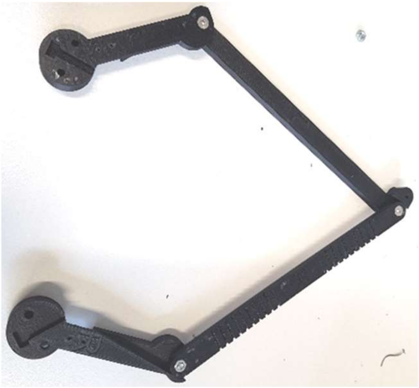

ii. **Install main body base**

10. Insert 1 M3 nut in each cavity of the main body as shown below.

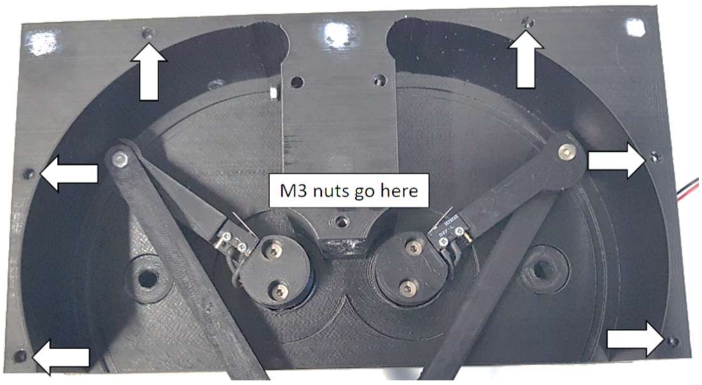


11. Insert 1 M3 nut in each cavity of the main body as shown below.

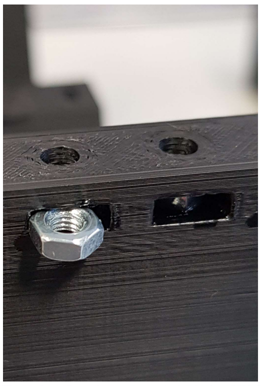

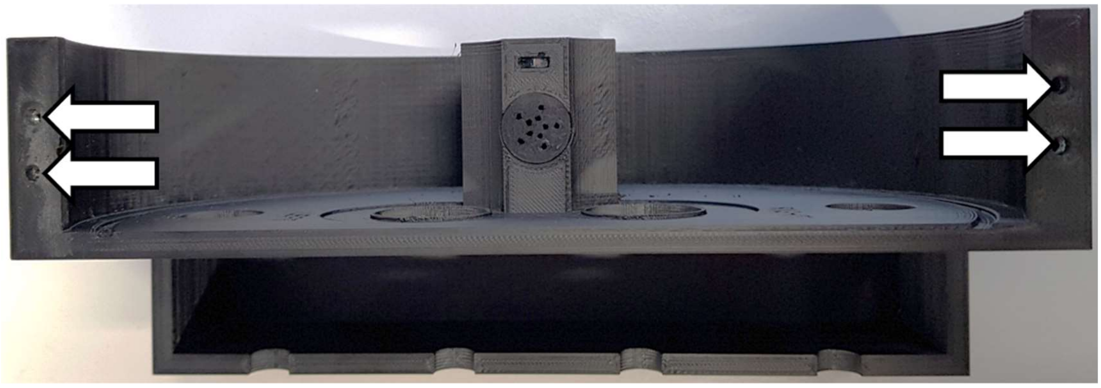

12. Fix the main body base to the main body using 6x 16mm M3 screws as shown below.

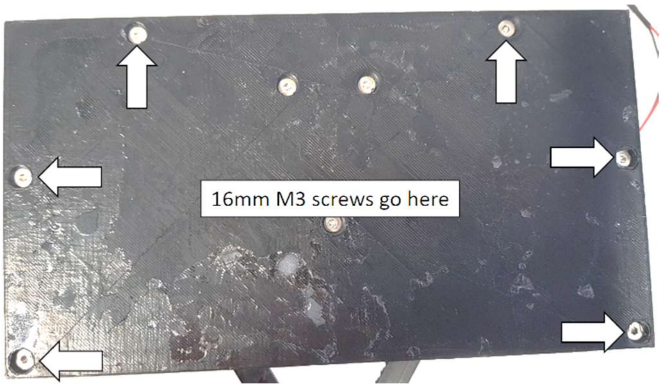


**b. Incubation Chamber**

**i. Assemble Base**

1. Insert 1x M3 nut into the horizontal cavities of the chamber body as shown below.

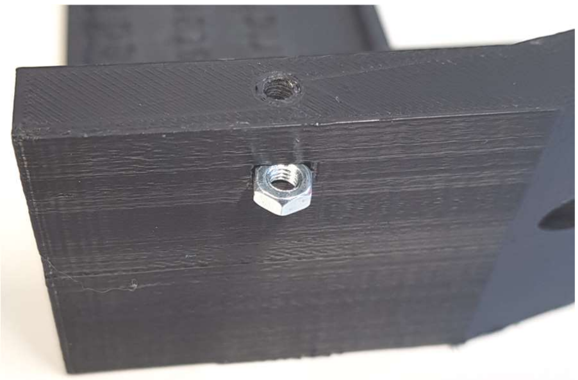

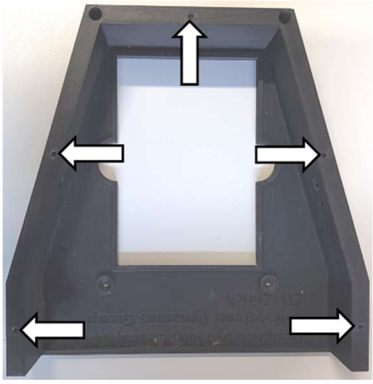

2. Place 1x 16mm M3 screw in each mounting cavity as shown below.

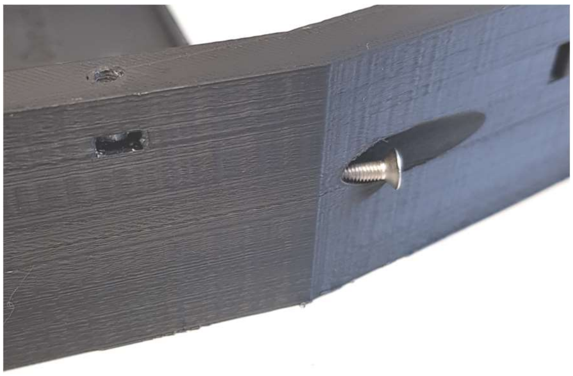

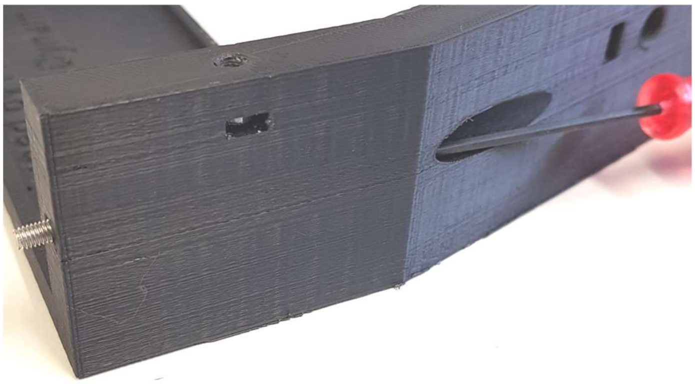

3. Screw gas connector into connection holes as shown below.

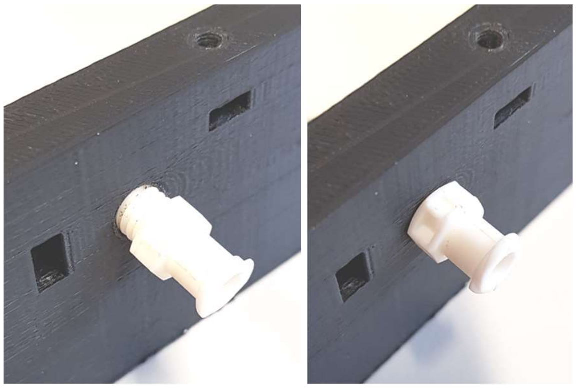

4. Attach Chamber lid to chamber body using 20mm M3 screws as shown below.

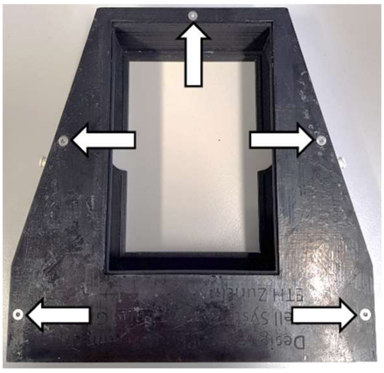

**c. Peristaltic Pumps**

**i. Assemble Peristaltic Pump Wheel**

a. **Supplies**

   i. **Wheel Bottom x 1**
   ii. **Wheel Top x 1**
   iii. **M3 Nut x 8**
   iv. **10mm M3 screw x 8**
   v. **8mm Bearing x 8**

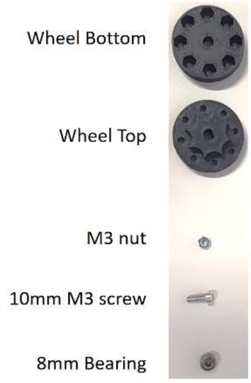


2. Place 10mm M3 screws in Wheel Top holes as shown below.

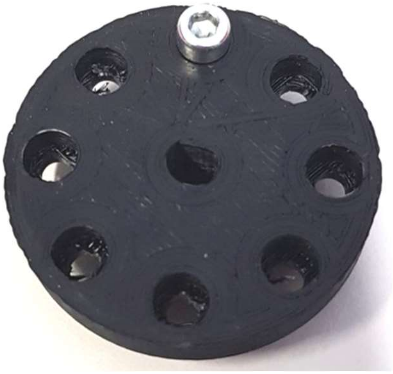

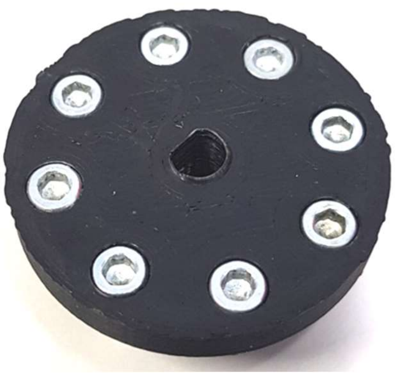

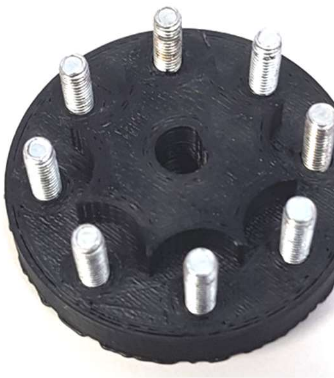

3. Place 8mm Bearings on 10mm M3 Screws as shown below.

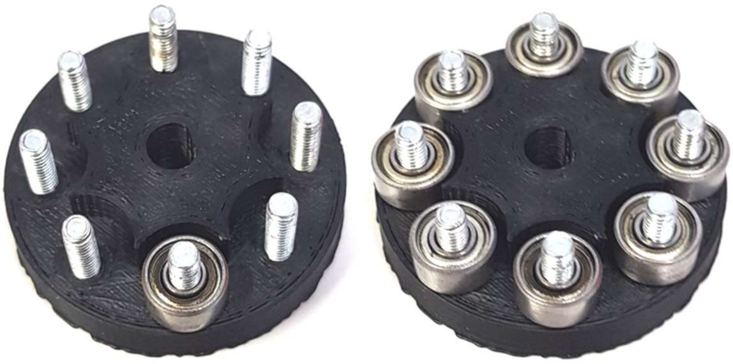

4. Place Wheel Bottom over Wheel Top and install M3 nuts as shown below.
    a. Note: After tightening all 10mm M3 Crews loosen each until the ball bearing spins freely.

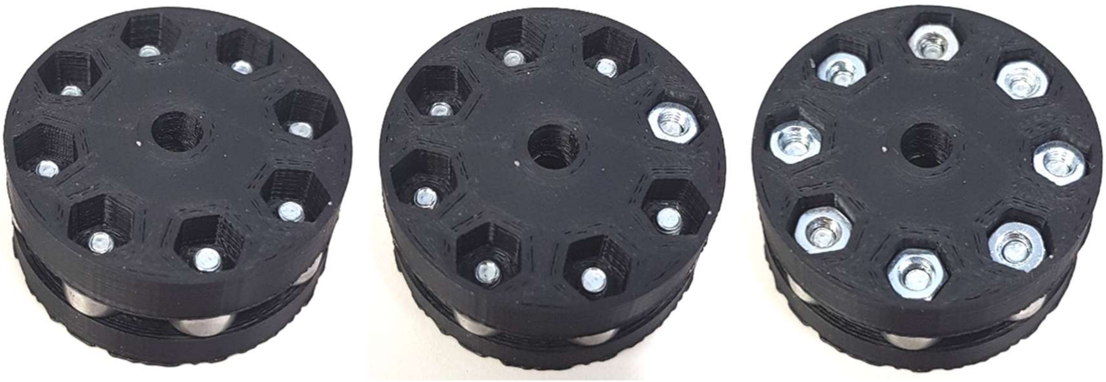

**ii. Assemble Pump Tubing**

**a. Supplies:**

**i.** Tygon LMT-55 Tubing

**ii.** 1/16” Barb Femal connector

**iii.** Scalpel

**iv.** 85mm Tube Cutting Jig

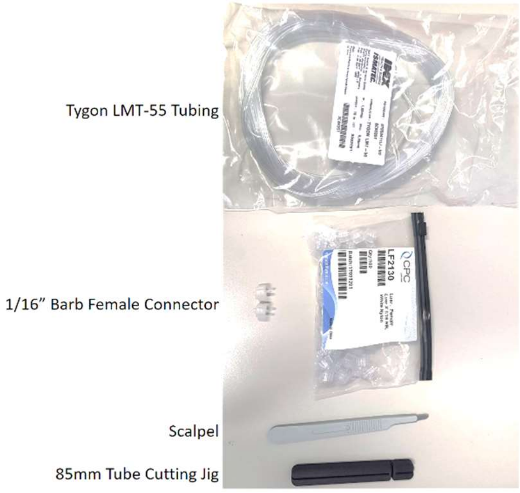


2. Cut Tygon LMT-55 Tubing into 85mm segment using the Tube Cutting Jig as shown below.

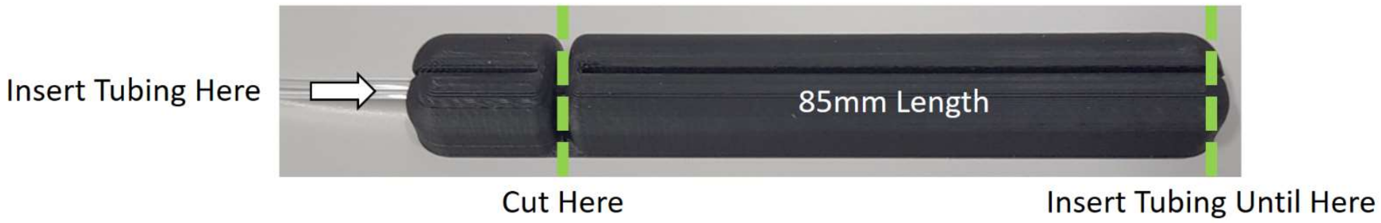


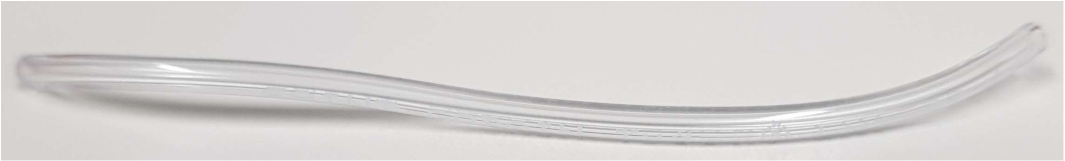

3. Attach 1/16” barb to each end of 85mm tube segment as shown below.

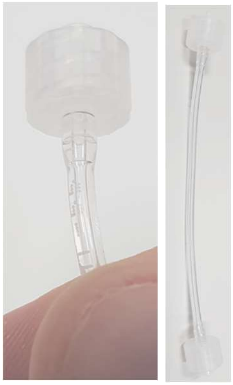

**iii. Assemble Peristaltic Pump Mount**

**a. Supplies**

**i. Nema 17 Stepper Motor x 1**
**ii. Peristaltic Pump Mount x 1**
**iii. Peristaltic Pump Bracket x 1**
**iv. 10mm M3 Screw x 2**
**v. 20mm M3 Screw x 2**

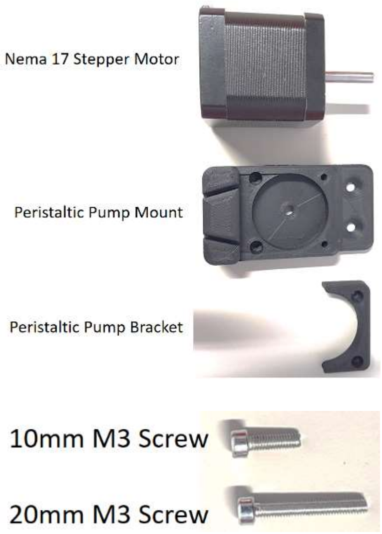


2. Place Peristaltic Pump Mount onto Nema 17 Stepper Motor and attach it with 10mm M3 Screws as shown below.

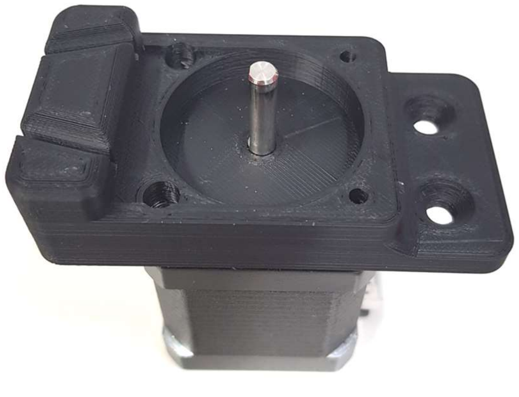

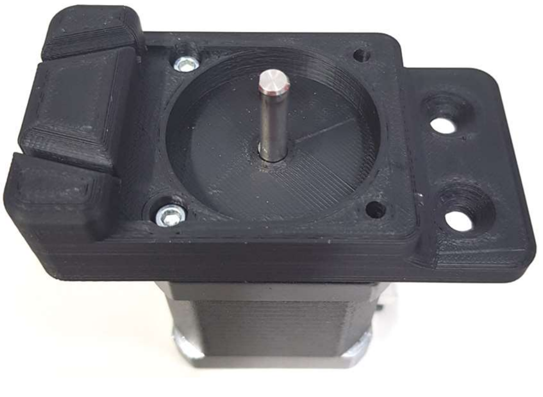

3. Place Assembled Wheel on Nema 17 Stepper motor shaft as shown below.
    a. Note: The flat edge of the wheel should match the flat edge of the Nema 17 Stepper Motor.

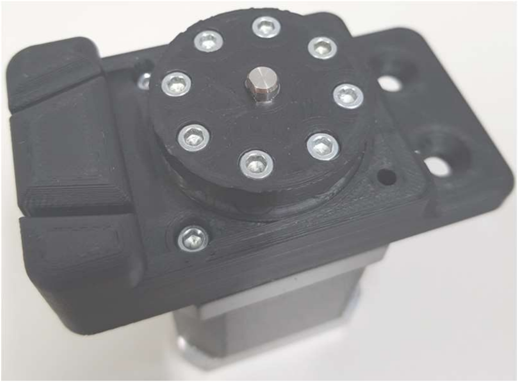

4. Place assembled Peristaltic Pump Tube in grooves as shown below.

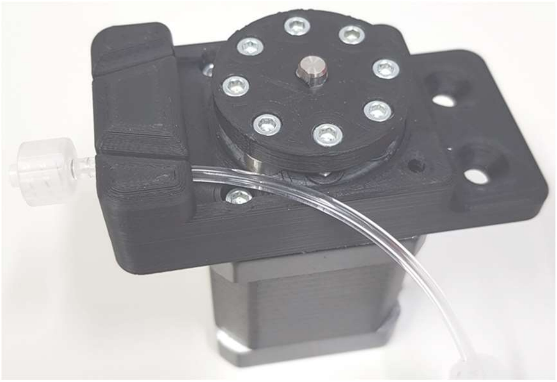

5. Stretch Peristaltic Pump Tube around the Wheel and through the second groove in the Peristaltic Pump Mount as shown below.

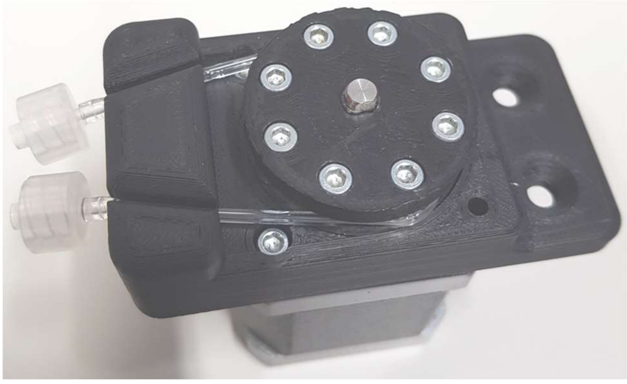

6. Place Peristaltic Pump bracket around tubing and wheel as shown below.

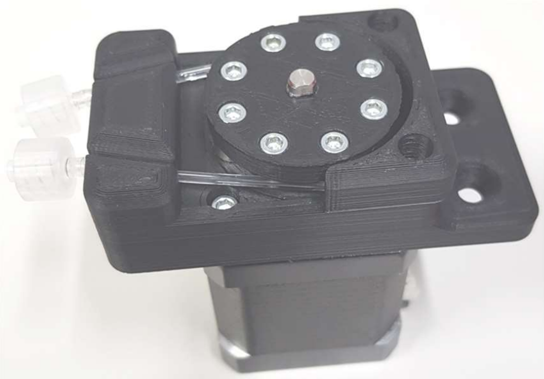

7. Attach Peristaltic Pump Bracket to Nema 17 Stepper Motor using 20mm M3 Screws as shown below.
    a. Note: You must apply pressure to the bracket to push it towards the wheel in order to align the holes for the screw.

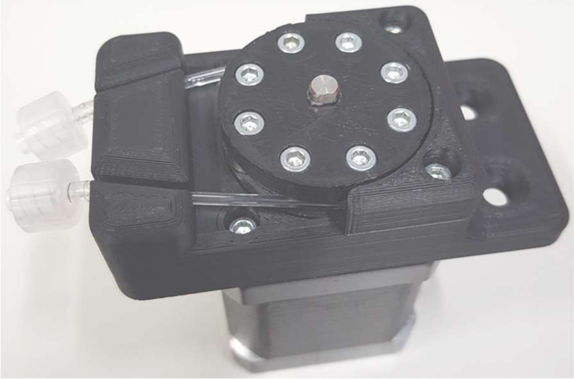

8. Repeat all steps for each Pump you wish to use.

**d. Control Box**

**i. Assemble Arduino Shield**

1. Gather 1x Arduino Mega, 5x 8 pin male header, 1x 6 pin male header, & 2x 18 pin male headers as shown below.

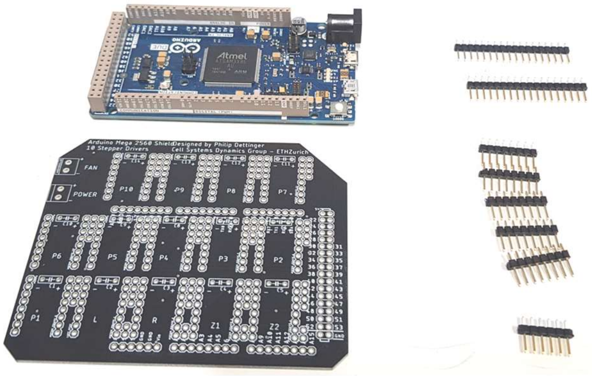

2. Insert the long end of each header into the female headers of the Arduino Mega as seen below.

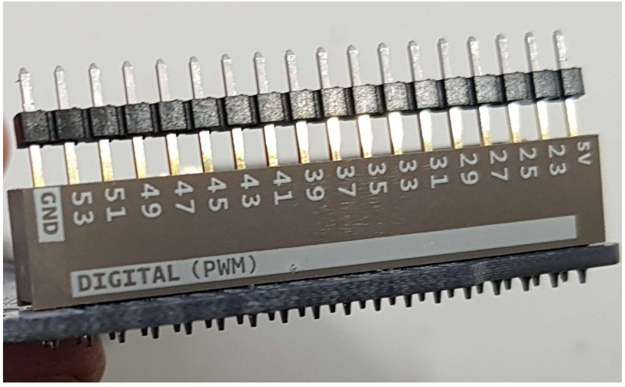


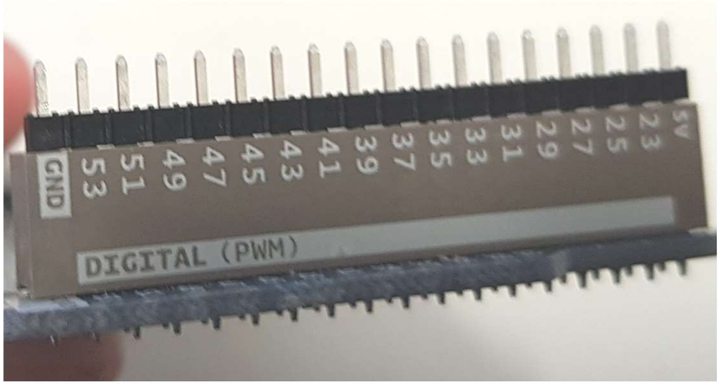


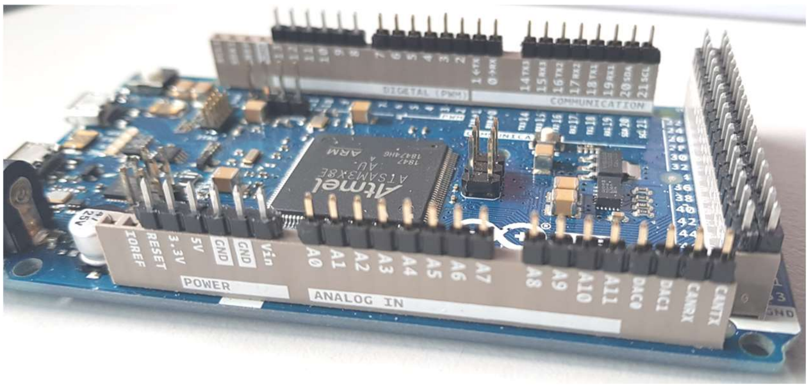


3. Place the Shield onto the Arduino Mega such that the male headers fit in the appropriate holes as shown below.

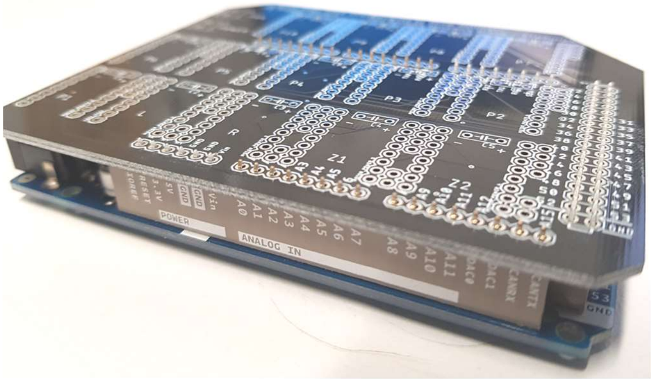

4. Solder each pin into place as shown below.

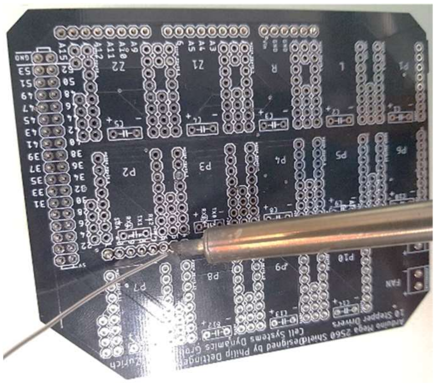

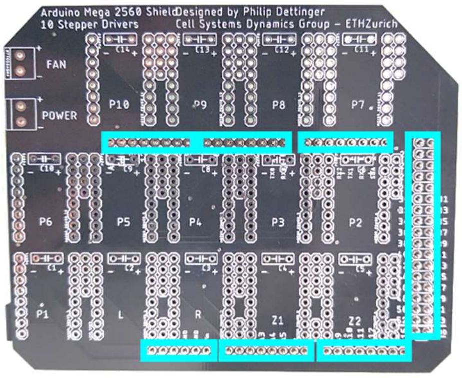


5. Place 8 pin female headers onto a stepper driver board as shown below.

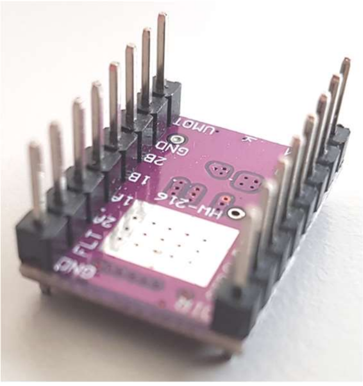

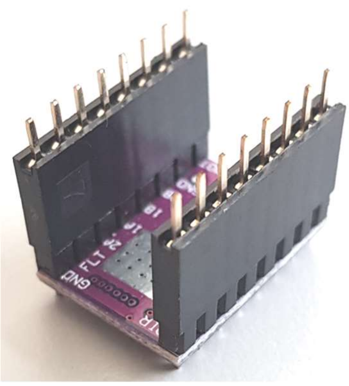

6. Place each stepper driver board into the shield holes then flip the board upside down using a clear plastic sheet to hold them in place as shown below.

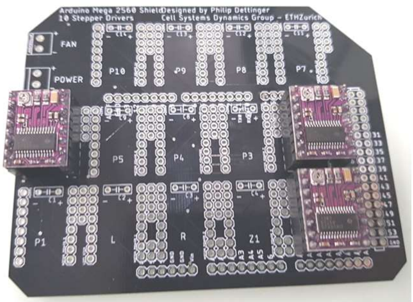

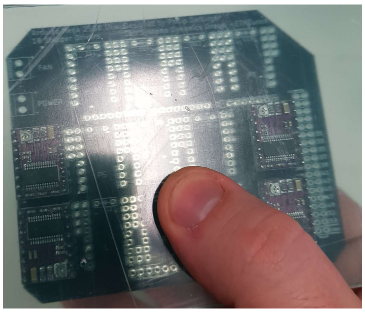

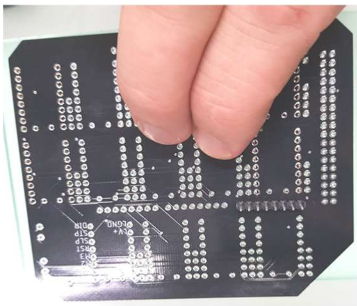

7. Solder stepper driver shields in place as shown below.

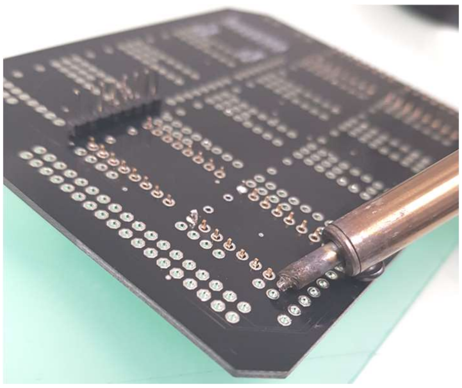

8. Repeat steps 6 & 7 for all stepper driver positions shown below.

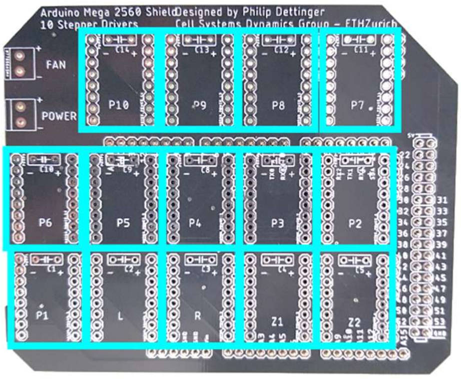


9. Place 4 pin & 2 pin female headers adjacent to stepper driver headers and hold them in place using 2 pin male headers then solder them to the bottom of the board as shown below.

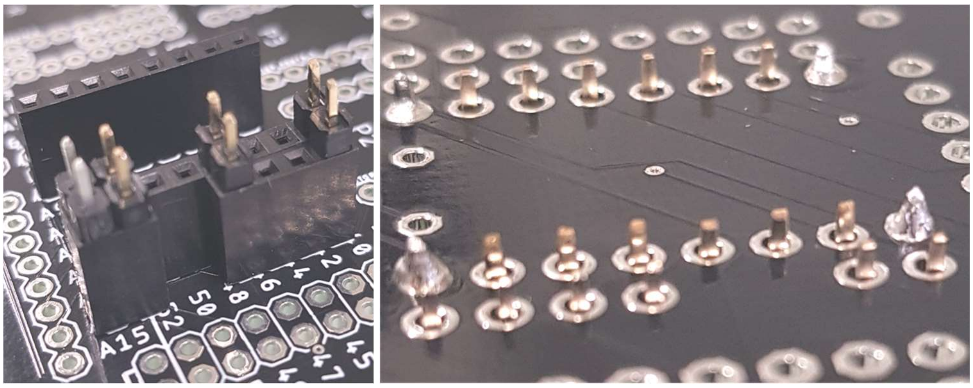

10. Repeat step 9 for all stepper driver boards shown below.
    a. Note: only 4 of the stepper driver boards require a 2pin female header.

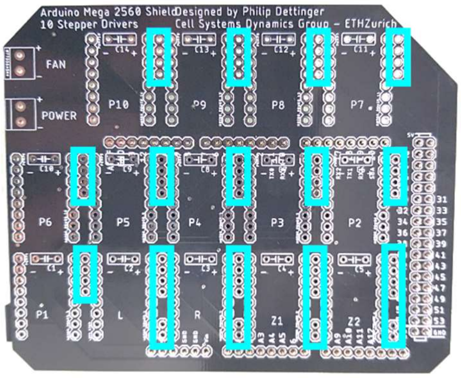


11. Place the power and fan terminal blocks into their appropriate holes. Bend their pins to hold them in place and solder them as shown below.

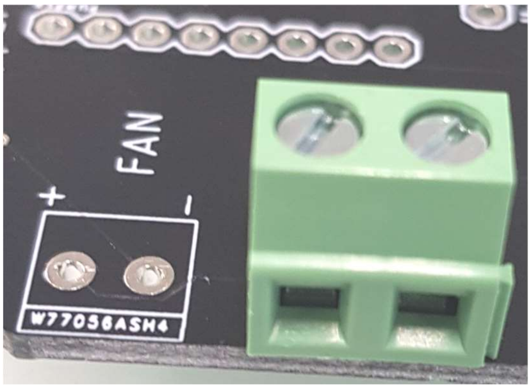

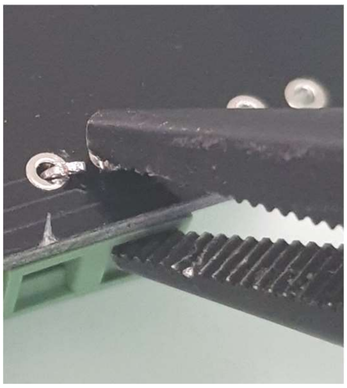
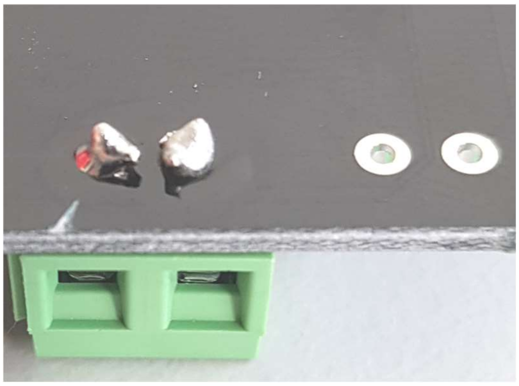

12. Place an electrolytic capacitor in its dedicated holes as shown below.
    a. Note: Electrolytic capacitors MUST be oriented correctly. If they are not they will boil and explode. PAY CAREFUL ATTENTION THAT THE NEGATIVE WIRE IS PLACED IN THE NEGATIVE HOLE AND THE POSITIVE WIRE IS PLACED IN THE POSITIVE HOLE AS SHOWN BELOW.

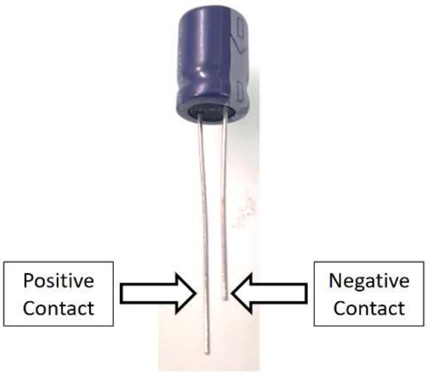


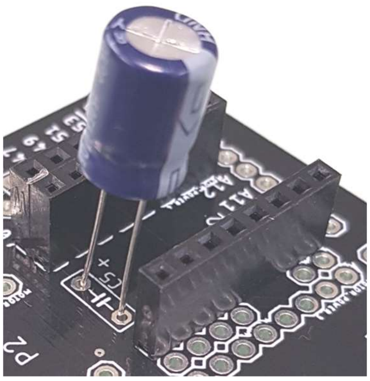

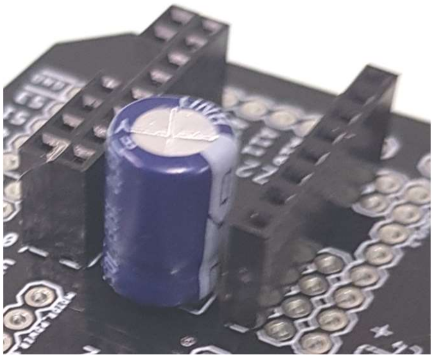

13. Bend the capacitor until it lays flush with the shield then bend the wires to hold it in place as shown below.

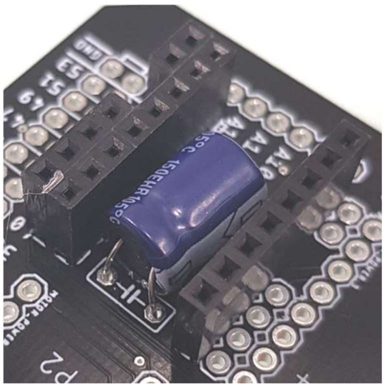

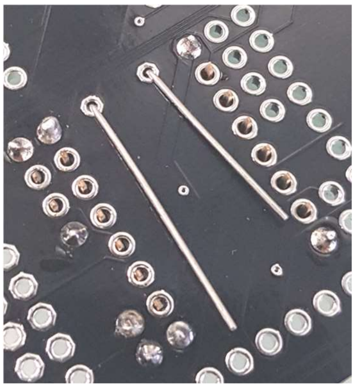

14. Solder the capacitor wires in place and clip away excess wire as shown below.

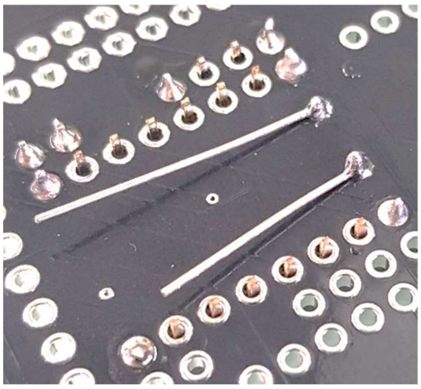

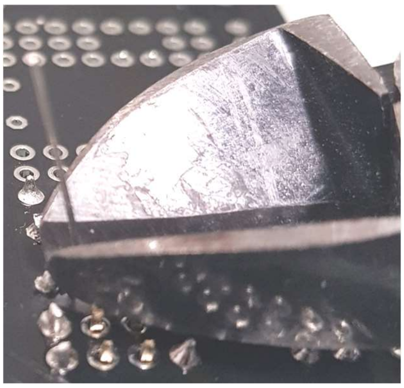

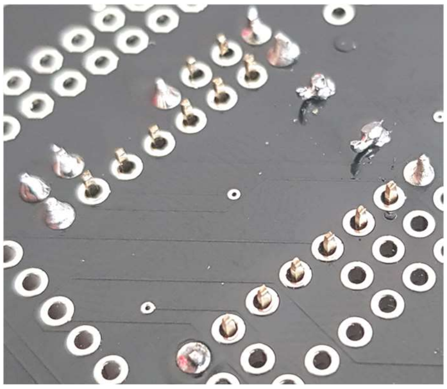

15. Repeat step 14 for all positions shown below.

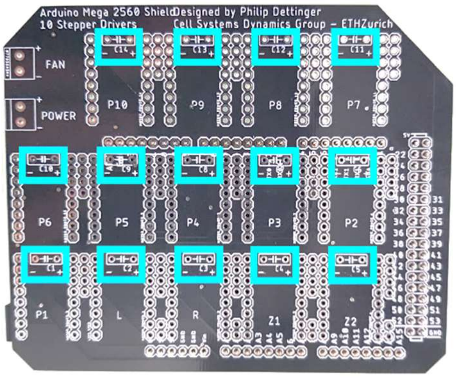


16. Your Arduino Shield should look like the photos shown below.

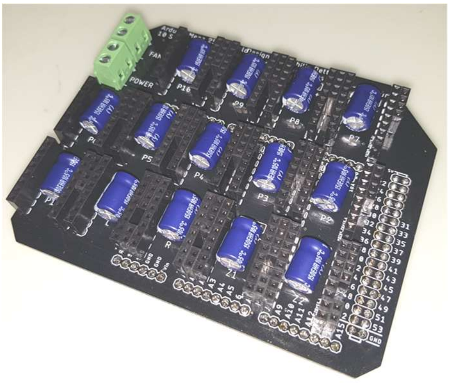

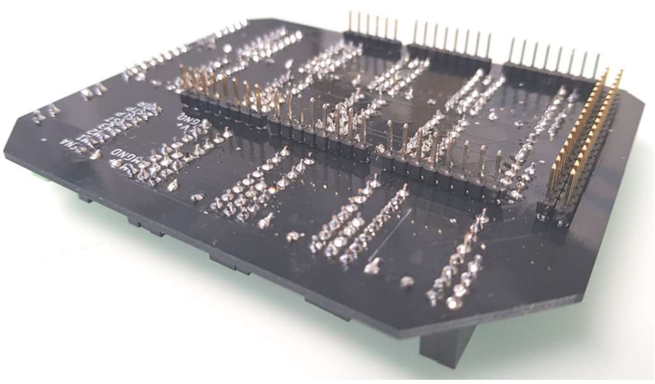

17. Place Arduino Mega on Arduino Plate as shown below.

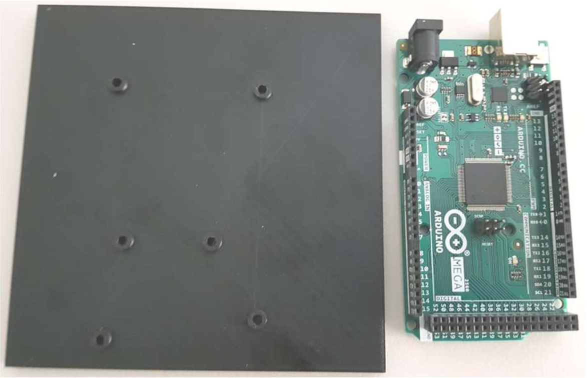 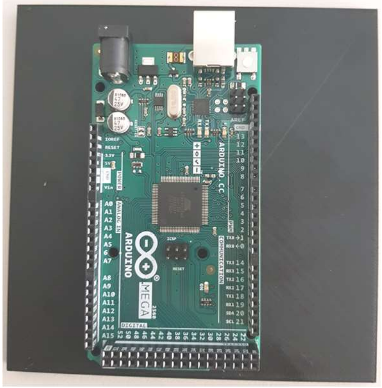

18. Attach the Arduino Mega to the Arduino Plate using 10mm M3 screws, M3 nuts, & glue as shown below.

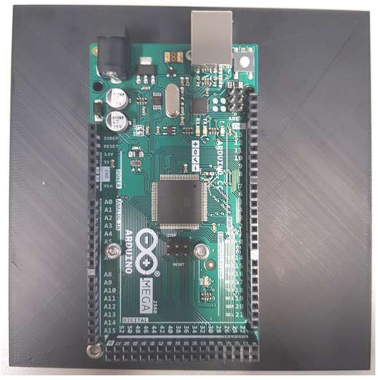 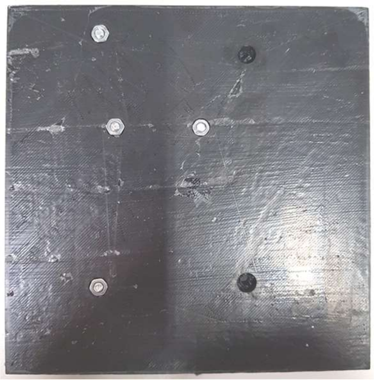 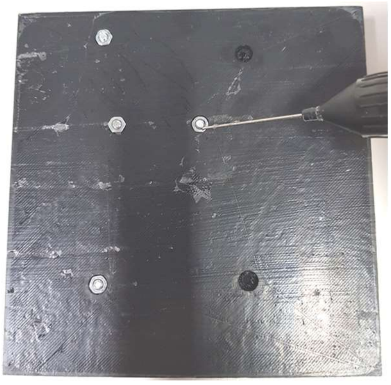

19. Place the Arduino Shield into the Arduino Mega.

**ii. Prepare Power Supply Box**

1. Screw 10mm M3 screws and M3 nuts in their respective places as shown below.

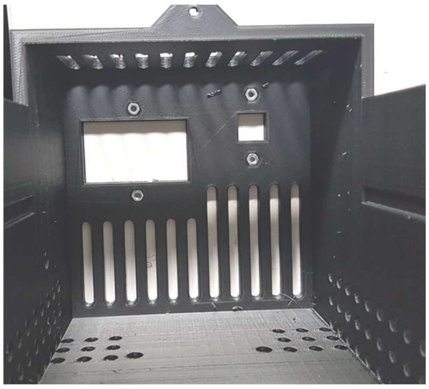 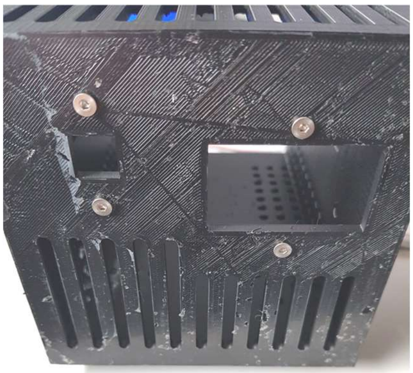

2. Glue the M3 nuts into place as shown below.

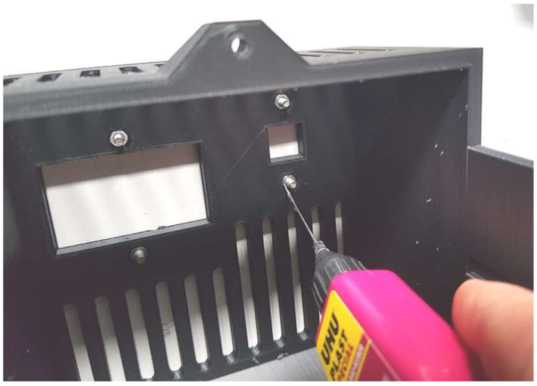

**iii. Assemble Power Supply**

1. Gather Power Switch, Connectors, Strippers, and Clippers as shown below.

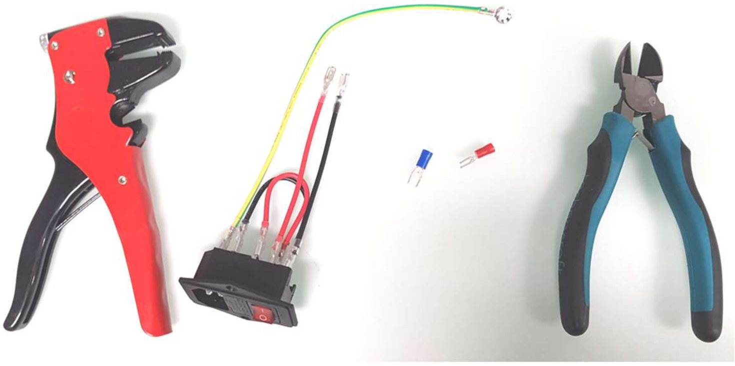

2. Clip off paddle connectors as shown below.

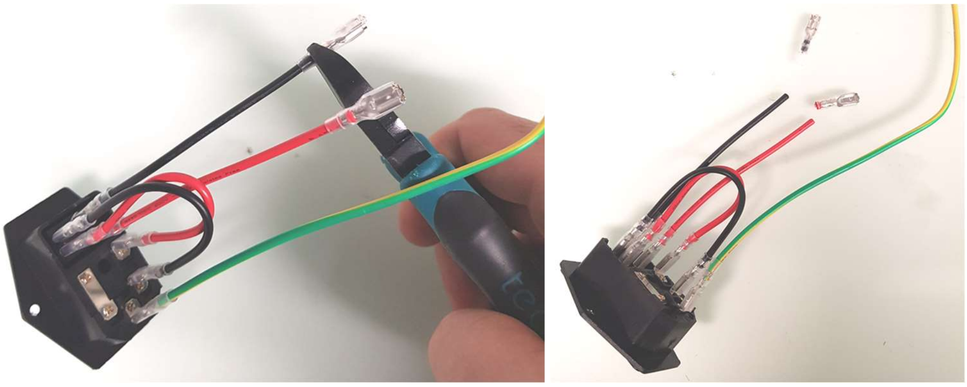

3. Strip wires 5mm from tip as shown below.

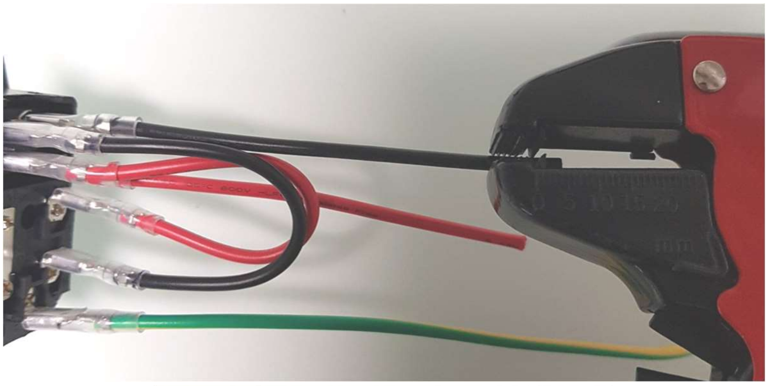

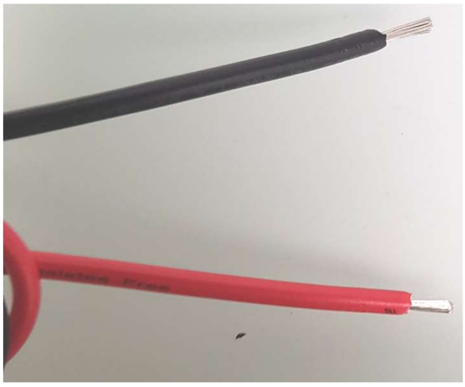

4. Insert bare wires into shovel connectors and solder as shown below.

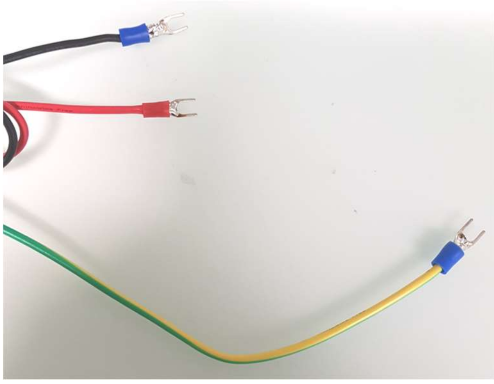

5. Insert power switch in the front panel of the box as shown below.

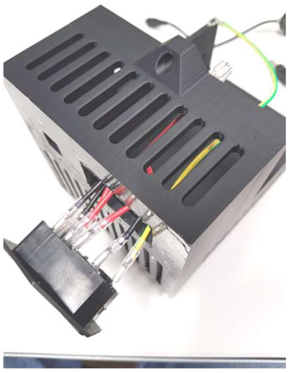

6. Loosen screws in the L, N, & GND positions of the power supply as shown below.

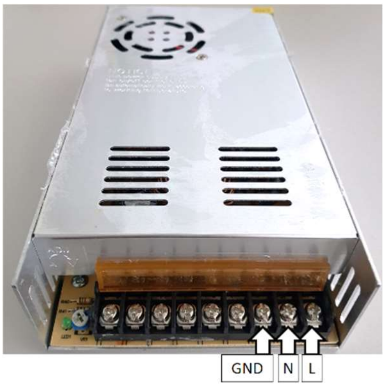


7. Place shovel connectors for L (Red), N (Black), and GND (Yellow/Green) in their respective locations and tighten their screws as shown below.

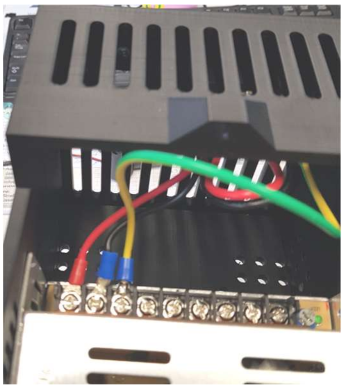

8. Connect V+ wire from the power supply to (+) power on the Arduino Shield and V- wire from the power supply to (-) power on the Arduino Shield as shown below.
9. Use 10mm M4 screws to connect the Power Supply to the box as shown below.
10. Connect V+ wire (Red) from the Fan to (+) fan on the Arduino Shield and V- wire (Black) from the Fan to (-) fan on the Arduino Shield as shown below.
11. Place Stepper Drivers into Driver ports according to your planned number of stepper drivers as shown below.
    a. NOTE: Pay close attention to their orientation of the Stepper drivers. The ENA pin must be in the upper right corner as shown below.
12. Thread Stepper Motor Cables through their holes in the Power Box and connect them to the Arduino Shield as shown below.
13. Repeat Step 11 for the limit switch cables as shown below.
14. Place the Arduino Plate in its slot as shown below.
15. Connect the Fan to the right half of the Power Box using 25mm M3 screws and M3 nuts as shown below.
16. Close the Power Box and fix the Power Supply using 10mm M4 screws as shown below.
17. Attach the handle to the box using 20mm M3 Screws and M3 nuts as shown below.

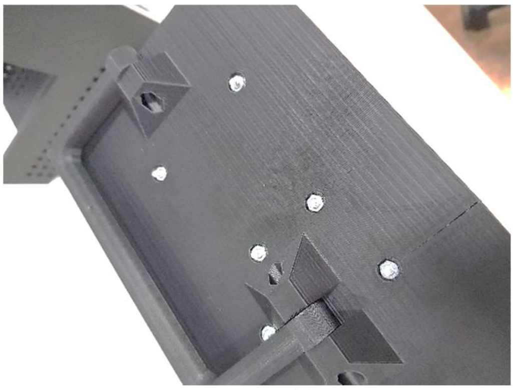

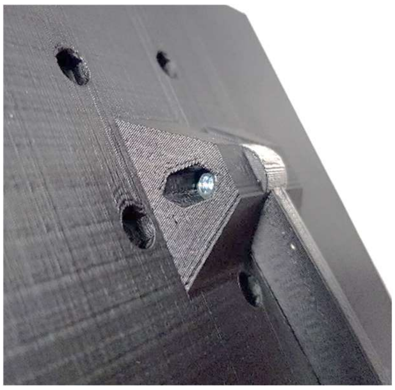

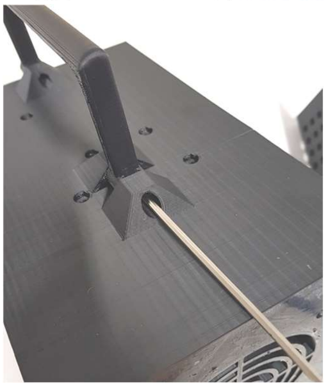

e. **Install Arduino Sketch**

   i. **Download Arduino IDE [https://www.arduino.cc/en/software](https://www.arduino.cc/en/software)**

   ii. **Connect to Arduino MEGA with USB cable.**

   iii. **Open Arduino IDE.**

   iv. **Open PHIL_14SM_210216 sketch.**

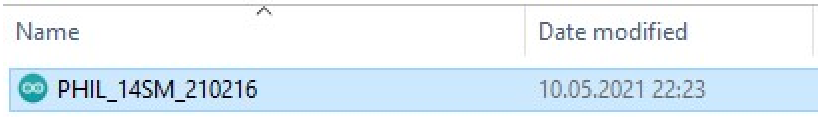


   v. **Select the correct port for connecting to the Arduino MEGA from Tools > Port**

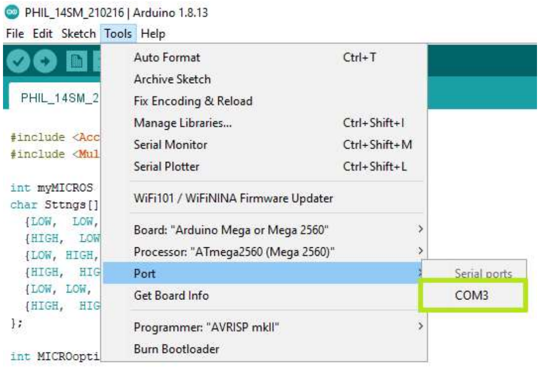


   vi. **Click "Upload."**

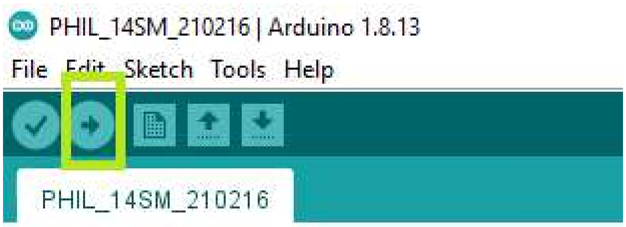


2. **Set Up PHIL**
    a. Place PHIL on desired surface.
    b. Fix PHIL in place with 4x 10mm M3 screws if placed on a microscope as shown below.

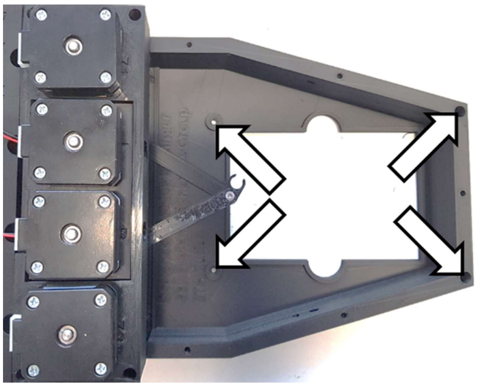

    c. Connect Power Control Box to stepper motors and limit switches as shown below.

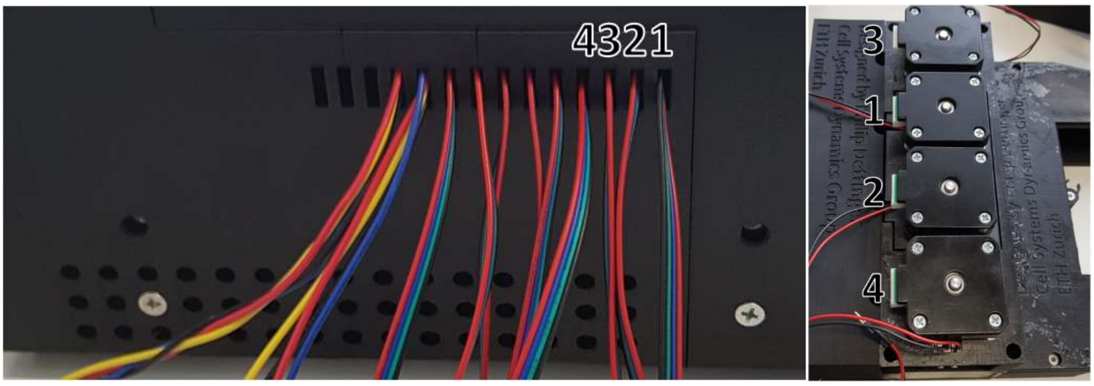


    d. Plug remaining stepper motor cables into peristaltic pumps as shown below.
        i. Note: All stepper motor cables must be plugged into a stepper motor to avoid damaging stepper motor drivers. Any stepper motor can be used for this purpose even if it is not to be utilized for pumping.

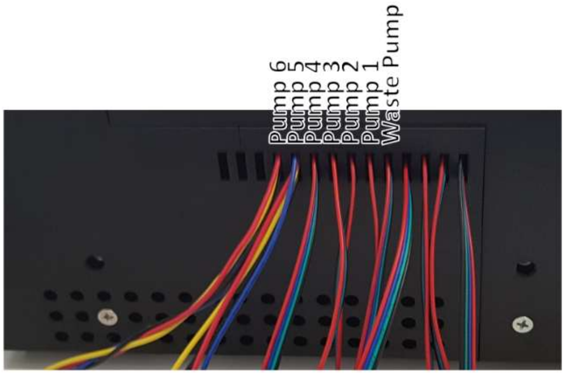

e. Plug power cord and USB into Power Control Box as shown below.

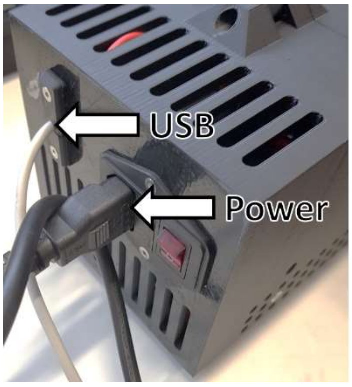


f. Plug power cord into power source and USB into the computer to be utilized for control.

3. **Appendix 1: Basic Soldering**
   a. **Useful resources:**
      i. https://www.instructables.com/How-to-solder/
      ii. https://www.youtube.com/watch?v=Zu3TYBs65FM
      iii. https://electronicsclub.info/soldering.htm
      iv. https://www.wikihow.com/Solder-Wires-Together
      v. https://uk.rs-online.com/web/generalDisplay.html?id=ideas-and-advice/how-to-solder-guide
   b. **Safety Precautions**
      i. **Soldering irons heat to >300°C. Never touch the tip with bare skin.**
      ii. **Soldering irons heat to >300°C. Never place them on combustible surfaces.**
      iii. **Soldering irons heat to >300°C. Hold them securely by the handle.**
      iv. **Solder often contains lead and produces smoke. Always solder in a well-ventilated area.**
      v. **Liquid solder is >300°C and can drip. Clear your soldering area of debris before soldering.**
      vi. **Liquid solder is >300°C and can drip. Wear the appropriate personal protective equipment (goggles, gloves, and a lab coat).**
      vii. **Soldering irons heat to >300°C. Always treat a soldering iron as though it were hot.**

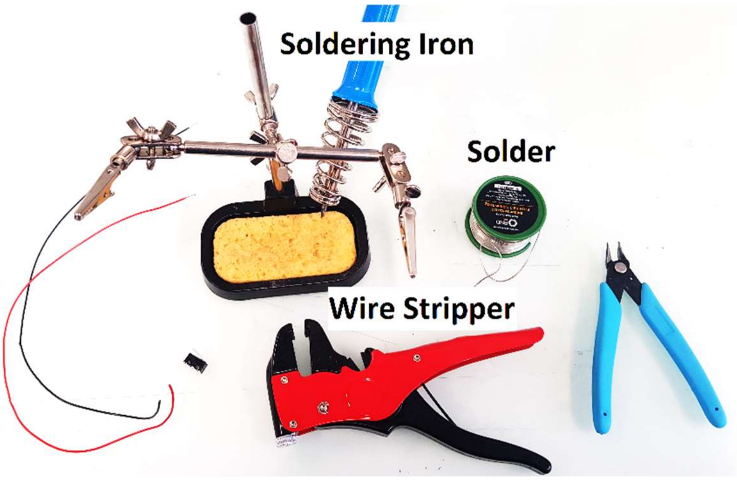


   c. **Step by Step**

i. Preheat your soldering iron by plugging it in to a power outlet.
ii. Check the temperature by touching the tip to a wet sponge.
iii. “Wet” the tip with solder as shown below.
    1. Note: If solder does not melt instantaneously, wait 1 minute and try again.

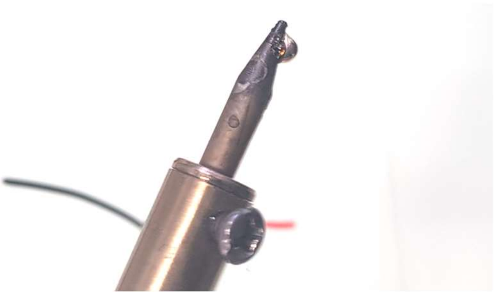

iv. Strip 5mm of wire insulation as shown below.

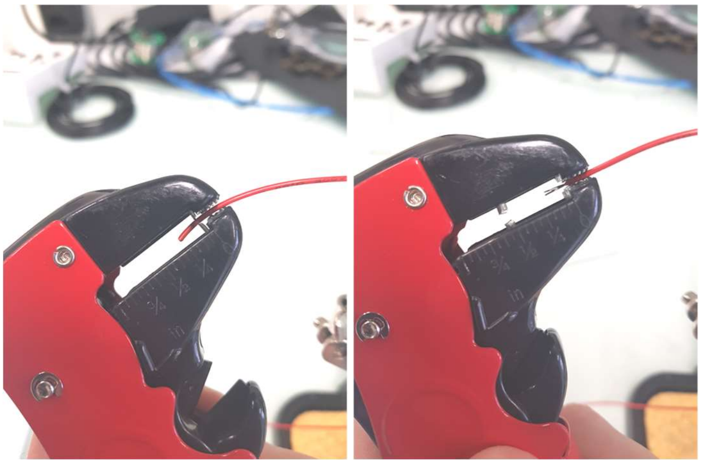

v. Splice the exposed wire with the joint as shown below.

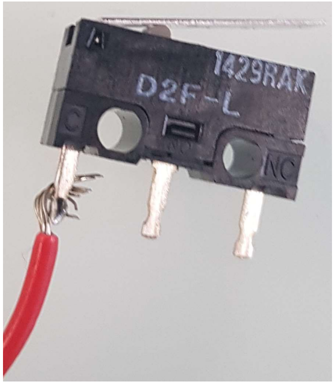


vi. Place your soldering iron tip to the spliced join and apply solder until all exposed metal has been covered and a secure joint is formed as shown below.

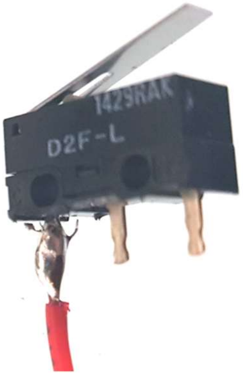


vii. Wipe your soldering iron on a wet sponge to clean excess solder and flux as shown below.

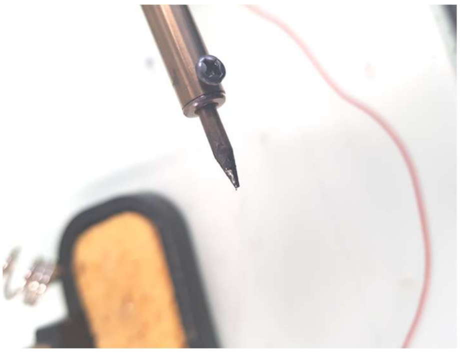
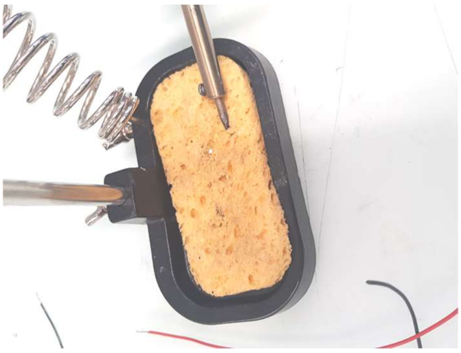
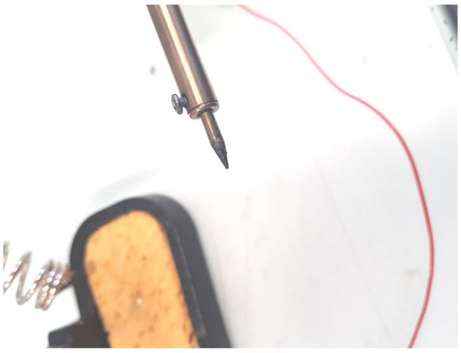

viii. Repeat as necessary.
